# VPS34 regulation in pulmonary arterial vascular smooth muscle cells in pulmonary hypertension

**DOI:** 10.64898/2026.09.28.752142

**Authors:** Anthony Natoli, Benjamin Reinhard, Logan Hallee, Huan Zhang, Kaylin Piza-Taylor, Felix Aung, Andrianna Poznahovska, Jason P. Gleghorn, Rachel C. Lucas, Maggie DiVita, Yuanjun Shen

**Author notes:** **Corresponding author**: Dr. Yuanjun Shen,. Department of Pharmaceutical Science, Binghamton University School of Pharmacy and Pharmaceutical Science, 96 Corliss Ave, Johnson City, NY, USA, 13790. **Author Contributions**: A.N. and Y.S. conceived and designed the study. A.N., B.R., K.P., H.Z., F.A., A.P., L.H., and Y.S. acquired the data. A.N., M.D., and Y.S. performed or verified the statistical analysis. L.H., F.A., and J.P.G. validated the protein-protein interaction prediction model. A.N., R.L., and Y.S. drafted the manuscript. M.D., R.L., and J.P.G. provided critical comments on the manuscript. All authors approved the final draft of the manuscript.

## Abstract

Pulmonary arterial hypertension (PAH) is characterized by the hyperproliferative phenotype of pulmonary arterial smooth muscle cells (PAVSMCs) and extensive extracellular matrix (ECM) remodeling. Vacuolar protein sorting 34 (VPS34), a Class III Phosphatidylinositol-3 Kinase (PI3K), regulates cell proliferation, but its specific function in pulmonary vascular remodeling remains poorly understood. We aim to determine the role of VPS34 activation and regulation in PAVSMC proliferation, matrix deposition, and the progression of PAH. First, we found a marked decrease in the inhibitory phosphorylation of VPS34 at Ser164 (P-S164-VPS34), indicative of constitutive VPS34 activation, in remodeled pulmonary arteries (PAs) from human PAH patients and male mice exposed to Sugen/Hypoxia (SuHx). In human PAH PAVSMCs, siRNA-mediated knockdown of VPS34 or pharmacological inhibition via the selective inhibitor SAR405 significantly blunted cell proliferation and reduced the production of fibronectin, a major component of the remodeled ECM. Mechanistically, we found that Akt activation in PAH PAVSMCs promotes VPS34 activation; Protein Phosphatase 1 Catalytic Subunit Alpha is a potential effector in VPS34 activation; and active VPS34 led to the activation of the mTOR/S6 signaling pathway with tuberous sclerosis complex 2 downregulation. Finally, *in vivo* treatment with SAR405 attenuated right ventricular systolic pressure, right ventricular hypertrophy, and pulmonary vascular remodeling in SuHx mice, while concurrently decreasing vascular fibronectin deposition. Our study demonstrates that VPS34 activation, driven by an Akt-mediated loss of the inhibitory P-S164, is crucial in PAVSMC proliferation and fibrotic vascular remodeling in PAH. Targeting the VPS34 axis represents a promising anti-remodeling therapeutic strategy for the treatment of PAH.

**Clinical Relevance:** *What is new?:*

1. This study identifies VPS34 activation via loss of the inhibitory Ser164 Phosphorylation as a previously unrecognized mechanism promoting pulmonary arterial smooth muscle cell (PAVSMC) hyperproliferation, pulmonary vascular remodeling, and pulmonary hypertension.
2. VPS34 activation increases PAVSMC-derived fibronectin production, highlighting the potential role of VPS34 in regulating extracellular matrix remodeling.
3. Pharmacological inhibition of VPS34 by SAR405 decreased PAH PAVSMC proliferation and pulmonary vascular remodeling.

*What are the clinical implications?:*

1. There is no FDA-approved VPS34 inhibitor, but chemicals or macromolecules targeting VPS34 activation, at least in PAVSMC, could be beneficial against pulmonary vascular remodeling and pulmonary hypertension progression.
2. Further understanding the regulation of extracellular matrix remodeling by VPS34 can also spark new therapies for pulmonary vascular diseases.

## Introduction

Pulmonary arterial hypertension (PAH) is a rapidly progressive cardiopulmonary disorder characterized by elevated pulmonary arterial pressure and right ventricular dysfunction. A hallmark feature of PAH is excessive proliferation of pulmonary artery (PA) resident cells, contributing to pulmonary vascular remodeling and increased vascular resistance^1^. Among these cellular changes, the hyperproliferation of PA vascular smooth muscle cells (PAVSMCs) plays a central role in driving pathological vascular remodeling.

Although the mechanisms underlying PAVSMC hyperproliferation in PAH are still not fully understood, increasing evidence suggests that the phosphatidylinositol 3-kinase (PI3K)/Akt/mammalian target of rapamycin (mTOR) signaling pathway plays a key regulatory role. For example, the activation of the p110 subunit of PI3K is required for PAVSMC proliferation and survival in PAH ^2,3^. Conversely, stabilization of tuberous sclerosis complex 2 (TSC2), a critical upstream inhibitor of mTOR complex 1 (mTORC1), induces apoptosis in PAH PAVSMCs ^4^. These findings suggest that dysregulation of the PI3K/Akt/mTOR axis contributes to aberrant PAVSMC growth; however, the full spectrum of its regulatory components and downstream effectors in PAH remains to be clarified.

PIK3C3, also known as vacuolar protein sorting 34 (VPS34), is the sole member of the Class III PI3K family and a pivotal mediator of phosphatidylinositol 3-phosphate (PtdIns3P) production. VPS34 orchestrates key cellular processes, including vesicle trafficking, autophagy, and lysosomal protein transport ^5,6^. Its activation depends on post-translational modifications (PTMs) and complex formation with regulatory proteins such as VPS15 ^7–9^, and activated VPS34 can stimulate mTOR signaling by destabilizing TSC2 ^10–12^. The inhibition of VPS34 through phosphorylation on Ser164 ^13^ and Thr159 ^14^ decreases cancer cell proliferation and promotes the death of human colorectal cancer stem cells ^15^ and breast cancer cells ^16^. Interestingly, SUMOylation of VPS34 has been implicated in promoting hyperproliferation of PAH PAVSMCs ^17^, suggesting that VPS34 may function as a cancer-like growth regulator in this disease context. However, the mechanisms governing VPS34 activation and function in PAH PAVSMCs are not well defined.

In this study, we investigated the status and regulation of VPS34 in PAVSMC proliferation and survival in the context of PAH. Our results demonstrate that VPS34 activation is Akt-dependent and required for the hyperproliferative phenotype of PAH PAVSMCs. These findings reveal a previously underexplored mechanism contributing to pulmonary vascular remodeling and highlight VPS34 as a potential therapeutic target in PAH.

## Methods

### Human Tissues and Cell Culture

Human lung tissues from healthy donors (control) and idiopathic PAH patients were provided by the Pulmonary Hypertension Breakthrough Initiative (PHBI). Cell characterization and maintenance were performed under the rigorous and well-established study protocols adopted by the PHBI. Briefly, PAVSMCs were characterized using antibodies against three smooth muscle-specific markers (smooth muscle α-actin [SMA], smooth muscle myosin heavy chain, and SM22; Cell Signaling Technology) and cell morphology. Primary cells (3-9 passages) from a minimum of three control and three PAH subjects were used for each experimental condition. Cells and tissues from de-identified human subjects were used (**Supplemental Table 1**). Cells were maintained at 37 °C in a humidified incubator with 5% CO_2_. For serum deprivation, cells were maintained for 24-48 hours in basal PromoCell Medium (PromoCell) supplemented with 0.1% bovine serum albumin (BSA) (Thermo Fisher Scientific). The cells were regularly tested for mycoplasma contamination. Immunohistochemical, Immunocytochemical, and Immunoblot Analyses information can be found in the **Supplemental Material**s.

### VPS34 knockdown

These procedures were performed as described previously^4,18^. The siRNAs were purchased from Dharmacon (PerkinElmer). siRNA-induced VPS34 knockdown was performed according to the manufacturer’s protocol.

### Animals

All animal procedures were performed under the protocols approved by the Animal Care and Use Committees at the University of California (Davis), University of Delaware, and Binghamton University (Protocol# 17019864, AUP# 1406, and Project# 936, respectively). Animal model establishment and data collection are described in the **Supplemental Materials**.

### Bioinformatic Analysis

The canonical human proteome was downloaded from UniProt (UP000005640) on February 2nd, 2026. To find a likely intermediate phosphatase between the anchor proteins VPS34 and AKT1, we screened the proteome with the Synthyra Proteome Prioritization Pipeline (SPP) against the question: Which human proteins were within two hops of VPS34 or AKT1 (called anchor proteins), shared their subcellular compartment, and were a phosphatase or associated with PAH? SPP used supervised protein language models (pLMs) to predict each human protein’s interaction partners, subcellular localization, gene-ontology terms, Enzyme Commission numbers, cofactors, UniProt keywords, and InterPro domains. Where UniProt carried the same annotation, it was scored by evidence source, with experimental observation ranking above curation by homology, which in turn ranked above electronic assertion; a computational annotation was kept when no experimental annotation existed for that category. Agreement between independent prediction methods raised confidence, while predictors sharing a backbone were counted once. Details about SPP-assisted annotation can be found in the **Supplemental Materials**.

### Statistical Analysis

Statistical comparisons between two groups were performed by the Mann-Whitney U test (non-parametric) or the Student’s T-test (parametric), and considered statistically significant when the p-value was less than 0.05. Statistical comparisons among three or more groups were performed by the Kruskal-Wallis test (non-parametric), followed by Dunn’s pairwise comparison, or the one-way ANOVA (parametric), followed by Bonferroni post-hoc analysis. Pearson correlation and regression analyses were also performed. No data points or animals were excluded from the analysis. To determine animal group sizes, Power calculations were performed. Statistical analysis was performed using R. All R code was initiated by ChatGPT and verified with the statistician.

## Results

### VPS34 activation is associated with pulmonary vascular remodeling and PAVSMC proliferation

VPS34 activity can be regulated through four major PTMs: phosphorylation, acetylation, ubiquitination, and SUMOylation ^6,9,14,17,19–21^. Specifically, VPS34 phosphorylation on Ser164 (P-S164-VPS34) inhibited VPS34 activity^13^. Our immunohistochemical (IHC) analysis detecting P-S164-VPS34 showed a significant and almost 50% decrease in PA VSM from PAH patients (PAH) compared to that from healthy donors (Contr.) (1.000±0.116 vs 0.541±0.058, Contr. vs PAH, p=0.0286) (**Figure 1 A,B**). Similarly, immunoblot (IB) analysis of human PAVSMC indicated that the ratio of P-S164-VPS34 to total VPS34 is significantly decreased by over 80% in those from PAH patients (1.000±0.331 vs 0.198±0.039, Contr. vs PAH, p=0.0079) (**Figure 1 C,D**). To determine the role of VPS34 in PAVSMC proliferation and survival, we next knocked down VPS34 expression by siRNA in PAH PAVSMC (**Figure 1E**). This led to a significant decrease in PAH PAVSMC proliferation by immunocytochemical (ICC) analysis detecting Ki67^+^ cells (1.000±0.000 vs 0.510±0.149, siContr vs siVPS34, p=0.0369) (**Figure 1F)**. Similarly, 48-hr incubation of VPS34 inhibitor SAR405 (5μM) with PAH PAVSMC significantly decreased cell proliferation (1.000±0.000 vs 0.568±0.127, Veh vs SAR405 5μM, p=0.0369) (**Figure 1 G,H**). In addition, we found that P-S164-VPS34 was significantly decreased in the PA endothelial layer in lung samples from PAH patients, suggesting VPS34 may play similar roles in pulmonary vascular endothelial regulation (1.000±0.027 vs 0.839±0.043, Contr. vs PAH, p=0.0286) (**Supplemental Figure 1**).

**Figure 1.**
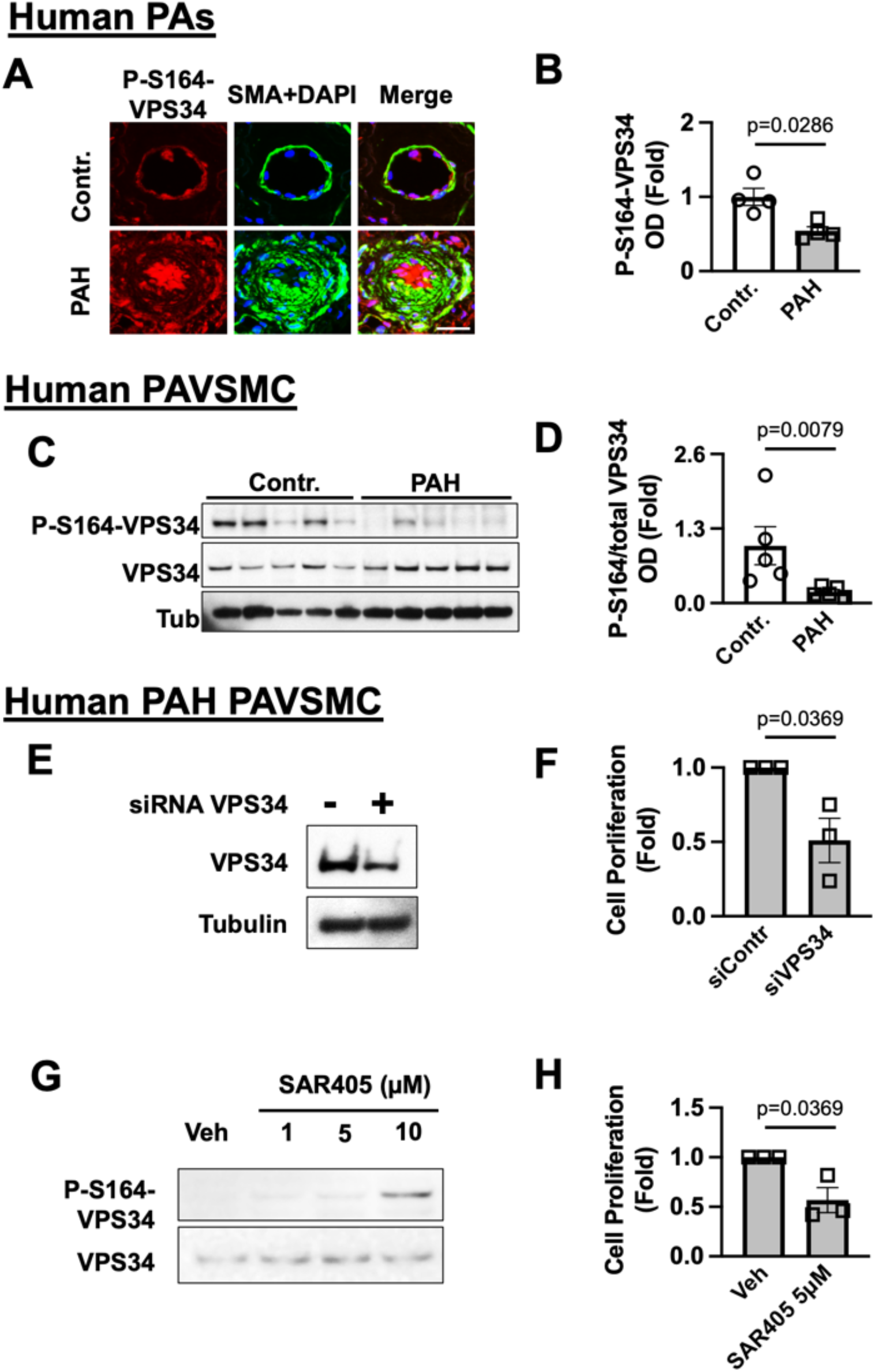
VPS34 is activated in human PAH PAVSMC. **A,B:** IHC analysis of human lung tissues to detect P-S164-VPS34 (red), SMA (green), and DAPI (blue). Bar=30 μm. Representative images (**A**) and optical density (OD) measurements of P-S164-VPS34 in SMA-positive regions of human small pulmonary arteries (**B**). **C,D**: Immunoblot analysis of PAVSMC from human non-diseased (control) and PAH lungs. Data are means±SE from n=5 subjects/group, analyzed by Mann-Whitney U test. Human PAH PAVSMCs were subjected to VPS34 knockdown by siRNA (**E**) or treated with SAR405 (VPS34 inhibitor) (**G**), confirmed by immunoblot analysis. **F, H:** Proliferation (Ki67) of human PAH PAVSMCs treated with SAR405. Data are % of Ki67-positive cells per total number of cells (DAPI); means±SE from n=3 subjects/group by Mann-Whitney U test.

### VPS34 is activated in PA VSM from male mice with SuHx-induced PH

To further understand the regulation of VPS34 *in vivo*, we induced experimental PH in male mice by SuHx, verified with right ventricular systolic pressures (RVSP, 26.579±0.828 mmHg vs 42.127±1.800 mmHg, Norm vs SuHx, p=0.00011), right ventricular hypertrophy (Fulton Index, 0.255±0.007 vs 0.452±0.032, Norm vs suHx, p=0.0012), and PA MT (1.000±0.077 vs 1.328±0.081, Norm vs SuHx, p=0.0167) (**Figure 2 A-D**). Similar to results from PAH patients, there was a decrease of P-S164-VPS34 (1.000±0.059 vs 0.668±0.036, Norm vs SuHx, p=0.0021) in VSM from remodeled small PA (<100 µm) in PH mice, which is negatively correlated with the increase in pulmonary vascular remodeling (P-S164-VPS34 ∼ PA MT, p=0.0014) (**Figure 2 E-G**).

**Figure 2.**
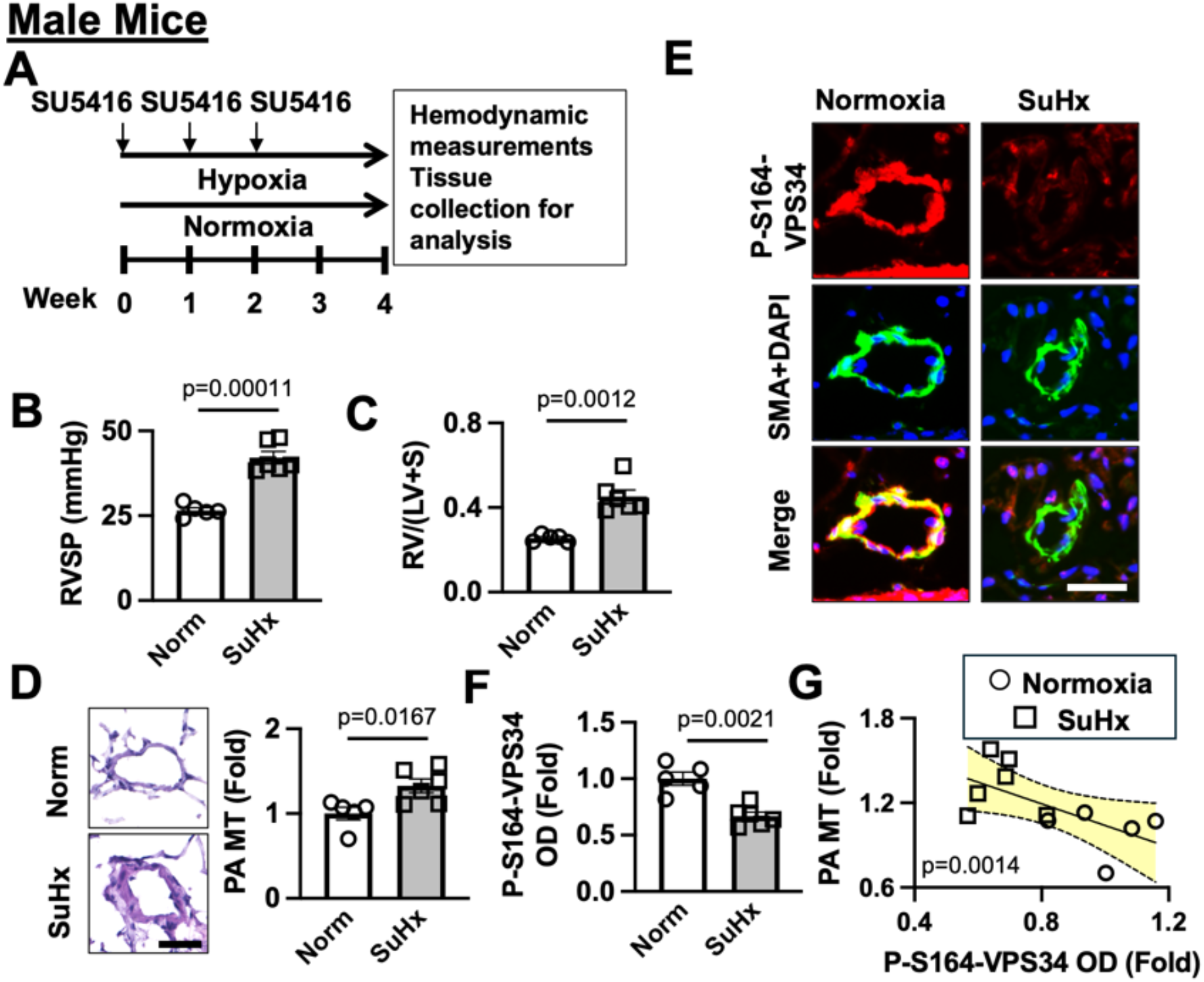
VPS34 is activated in PA SM in SuHx-induced PH in mice. **A.** Experimental Scheme: male 6-8 w.o. mice received SU5416 injections once a week for three weeks and were maintained under hypoxia for four weeks. Controls were same-age male mice kept under normoxia for 4 weeks. **B,C**. RVSP (**B**) and Fulton Index (RV/(LV+S)) (**C**) were measured on Day 28 of the experiment. **D**. H&E staining and PA MT of small pulmonary arteries: bar=50 μm; images are representative of 5 (Normoxia) or 6 (SuHx) mice/group, 12 pulmonary arteries/mouse; analyzed by Student’s T-test. **E,F**: IHC analysis of mouse lung tissues to detect P-S164-VPS34 (red), SMA (green), and DAPI (blue). Bar=30 μm. Images are representative of 5 (Normoxia) or 6 (SuHx) mice/group, 12 pulmonary arteries/mouse (**E**). OD values were calculated from 12 areas/pulmonary artery by Student’s T-test (**F**). G. Correlation analysis between PA MT and P-S164-VPS34 OD values by Pearson’s correlation.

### VPS34 activation in PAH PAVSMC is likely associated with Akt and protein phosphatases

It is known that PAVSMC hyperproliferation is associated with Akt activation^4,18,22^, which is also largely involved in PI3K pathway regulation^2,3,23,24^. We then tested whether Akt is an upstream regulator in VPS34 activation in PAVSMC. First, we incubated human PAH PAVSMC with vehicle (DMSO) or 1 and 5 µM Akt Inhibitor VIII (AKTi VIII) for 48 hours. The IB analysis showed a dose-dependent increase in P-S164-VPS34 in human PAH PAVSMC, suggesting that AKTi VIII promotes Ser164-phosphorylation-dependent VPS34 inactivation (1.000±0.000 vs 13.902±5.952 vs 18.939±8.661, AKTi VIII 0μM vs 1μM vs 5μM) (**Figure 3 A,B**).

**Figure 3.**
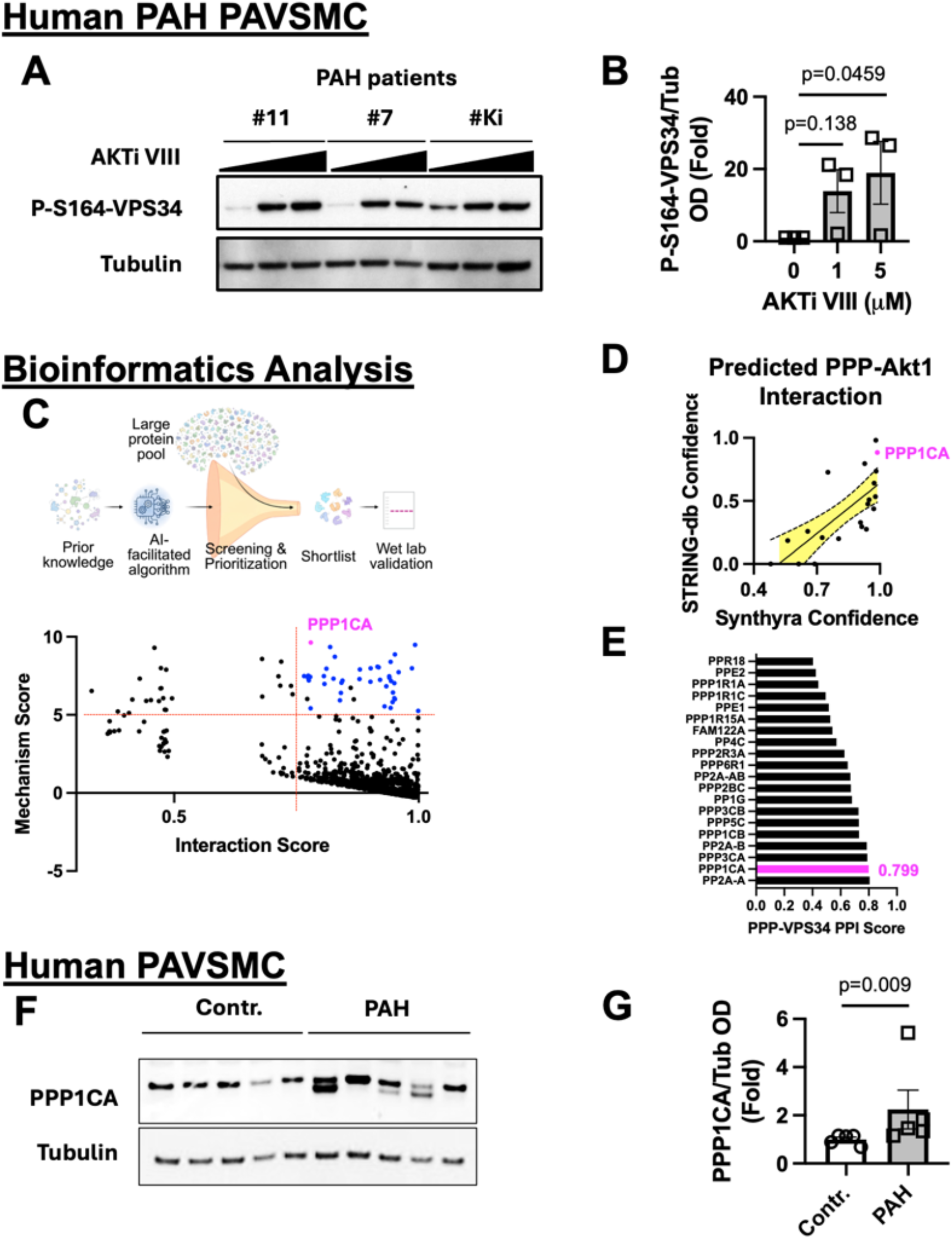
VPS34 activation in human PAH PAVMSC is regulated by AKT and potentially PPP1CA. **A,B:** Immunoblot analysis of human PAH PAVSMC treated with diluent (0), 1, and 5 μM AKTi VIII for 48hr to detect S164-VPS34 phosphorylation rates. Data are means±SE from 3 subjects/group, analyzed by Mann-Whitney U Test. **C**. Synthyra Proteome Prioritization Pipeline (SPP). The pipeline prediction is plotted as Mechanism Score vs. Interaction Score. 42 proteins with high mechanism score (>5) and interaction score (>0.75) are found in the top-right corner. PPP1CA is marked in pink. Other protein phosphatases are marked in blue. (**D,E**) STRING-db and SPP were used to analyze interaction scores between commonly reported PPPs and Akt or VPS34. For STRING-db, no reported association is shown as 0, while higher association scores approach 1. STRING-db-reported scores were divided by 100. **D**. STRING-db-reported scores were plotted against SPP-predicted interaction scores, and the correlation analysis was performed using Pearson’s correlation. PPP1CA is marked in pink. **E**. Commonly reported PPPs with their predicted interaction scores with VPS34. **F,G**: Immunoblot (IB) analysis detecting PPP1CA and Tubulin in human PAVSMCs from healthy donors (Control) and PAH patients. IB results were analyzed using ImageJ. Data are means±SE from 5 subjects/group, analyzed by Student’s T-test.

Akt is known to function as a kinase. To understand why Akt can promote the transition of inactive P-S164-VPS34 into active dephosphorylated VPS34, we hypothesized that one or multiple phosphatases may be involved in this regulation. We used SPP to nominate candidate phosphatases with predicted functional proximity to Akt1 and VPS34, and a potential role in PH progression. After screening and prioritization, we identified 42 phosphatases (**Figure 3C**), 39 of which are protein phosphatases (PPPs, in Blue). Therefore, we specifically chose PPPs to further test, as they were also previously reported to be responsible for cancer cell proliferation. Published data on interactions between PPPs and VPS34 are very limited; typically, STRING-db analysis returned most predictions as unknown (interaction score = 0). STRING-db and SPP showed concordant rankings (p=0.0003) for Akt1 and 20 commonly reported PPPs, most of which are already documented in the literature (**Figure 3D**). For VPS34-PPP pairs, which are largely undocumented, STRING-db returned no signal for nearly all pairs, while SPP generated a ranked candidate list (**Figure 3E**); because no ground truth exists for these pairs, we treated this ranking as a prioritization tool rather than a validated prediction. With these bioinformatic results, we found that PPP1CA was prioritized across all three analyses (in Pink, **Figure 3C-E**). Therefore, we tested the expression of PPP1CA in PAVSMC by IB, which showed that PPP1CA was significantly increased in lysates from PAH PAVSMC compared to those from control cells (1.000±0.093 vs 2.243±0.803, Contr. vs PAH, p=0.009) (**Figure 3 F,G**). However, more work is needed to elucidate whether PPP1CA mediates Akt-dependent VPS34 activation. The convergence of an unbiased proteome-wide nomination, elevated PPP1CA abundance in PAH PAVSMC, and PPP1CA’s prior reported roles in proliferation and Akt-associated signaling collectively support PPP1CA as a biologically plausible candidate warranting direct mechanistic testing (e.g., co-immunoprecipitation, siRNA knockdown).

### VPS34 activation regulates PAVSMC fibronectin production

Pulmonary vascular remodeling in PAH is known to be associated with extracellular matrix (ECM) remodeling^4,18^. Here, we tested whether VPS34 regulates the production of fibronectin (FN), an important component of the ECM, in PAVSMC. As described above, we used IHC to measure the level of FN in PA VSM from male mice maintained in normoxia or exposed to SuHx, with or without the administration of SAR405 (**Figure 4A**). As expected, compared to mice maintained in normoxia, FN levels were significantly increased in PA VSM from mice exposed to SuHx (1.000±0.042 vs 1.360±0.072, Norm vs SuHx+Veh, p=0.00167). Two-week administration of the VPS34 inhibitor SAR405 (10 mg/kg/day) in male mice exposed to SuHx significantly decreased FN levels in PA VSM (1.360±0.072 vs 1.115±0.045, SuHx+Veh vs SuHx+SAR405, p=0.0236) (**Figure 4B**). Importantly, FN levels in PA VSM are inversely correlated with P-S164-VPS34 (P-S164-VPS34 ∼ FN, p=0.00299) (**Figure 4C**).

**Figure 4.**
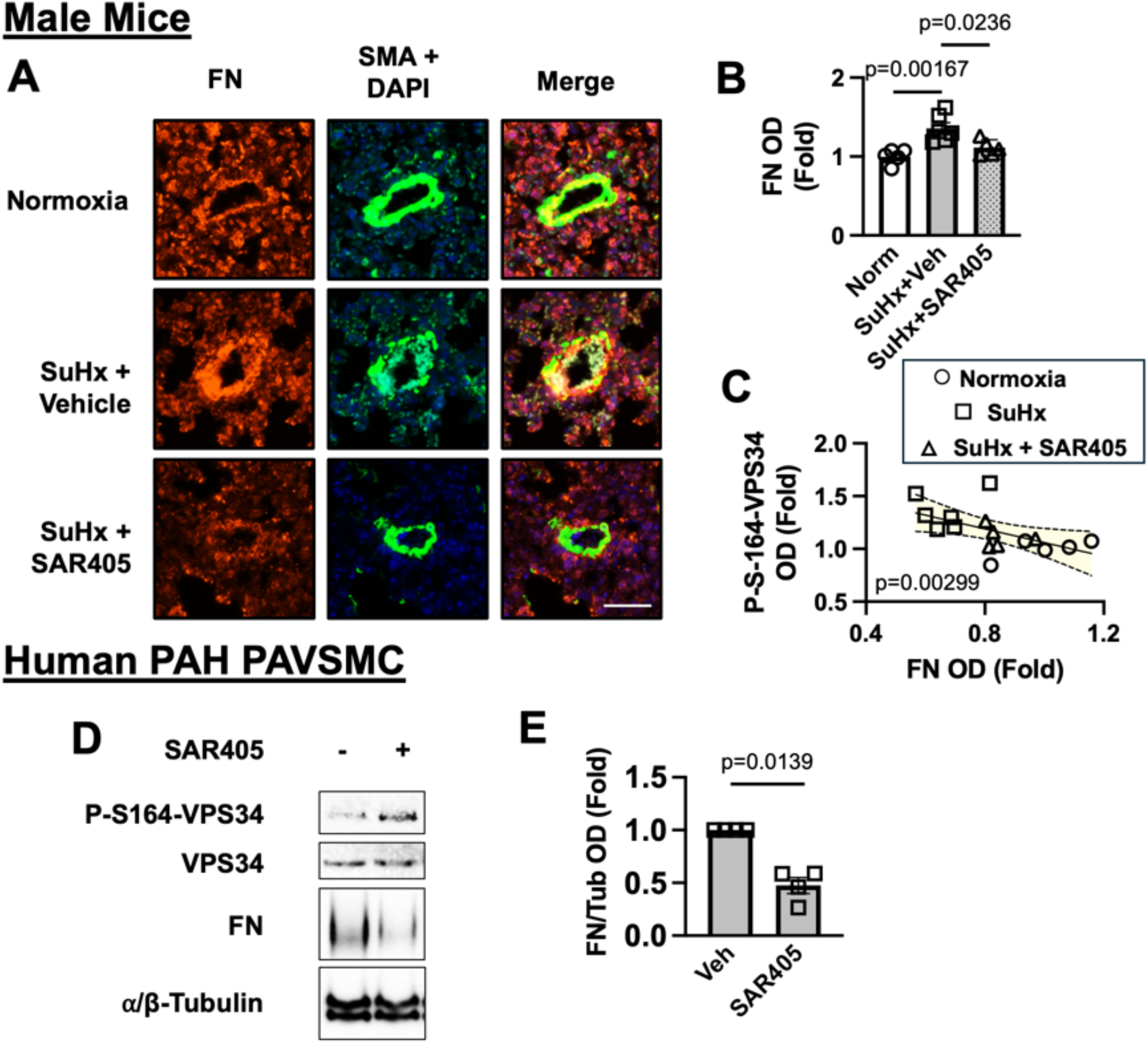
VPS34 regulates fibronectin (FN) in PH mice and PAH PAVSMC. **A,B:** IHC analysis of mouse lung tissues to detect FN (red), SMA (green), and DAPI (blue). Bar=50 μm. Images are representative of 5 (Normoxia), 6 (SuHx+Vehicle), and 5 (SuHx+SAR405) mice/group, 12 pulmonary arteries/mouse. OD values were calculated from 12 areas/pulmonary artery; statistical analysis was performed by ANOVA, followed by the Bonferroni post hoc test. **C**.Correlation analysis between FN and P-S164-Vps34 OD values using Pearson’s correlation. **D,E**: Immunoblot analysis of human PAH PAVSMC treated with SAR405, detecting FN and Tubulin. Data are means±SE from n=4 subjects/group, analyzed by Student’s T-test

To verify the *in vivo* results, PAH PAVSMCs were treated with SAR405 (10 μM) for 48 hours. IB analysis showed a significant decrease in FN (1.000±0.000 vs 0.476±0.076, Veh vs SAR405, p=0.0139), along with restored VPS34 phosphorylation, in PAH PAVSMCs treated with SAR405, further indicating that VPS34 is an upstream regulator of FN (**Figure 4 D,E**).

### VPS34 activation in PAH PAVSMC regulates VPS15 and TSC2 protein levels and mTOR activation

Next, we investigated the downstream effects of VPS34 activation in PAH PAVSMC. IB analysis was performed to detect the protein levels of VPS15 (a structural component of the VPS34 complex), TSC2 (the upstream negative regulator of mTORC1), and the phosphorylation of S6 (an indicator of mTOR activation)^4,6^ upon the knockdown of VPS34 by siRNA in PAH PAVSMCs. We found that VPS34 knockdown led to a significant decrease in VPS15 protein level (1.000±0.000 vs 0.873±0.104, vs siContr vs siVPS34, p=0.0369), a significant increase in TSC2 protein level (1.000±0.000 vs 1.455±0.329, vs siContr vs siVPS34, p=0.0369), and a significant decrease in S6 phosphorylation (1.000±0.000 vs 0.497±0.257, vs siContr vs siVPS34, p=0.0369) (**Figure 5 A-D**), demonstrating that VPS34 regulates TSC2 and, consequently, mTOR activation.

**Figure 5.**
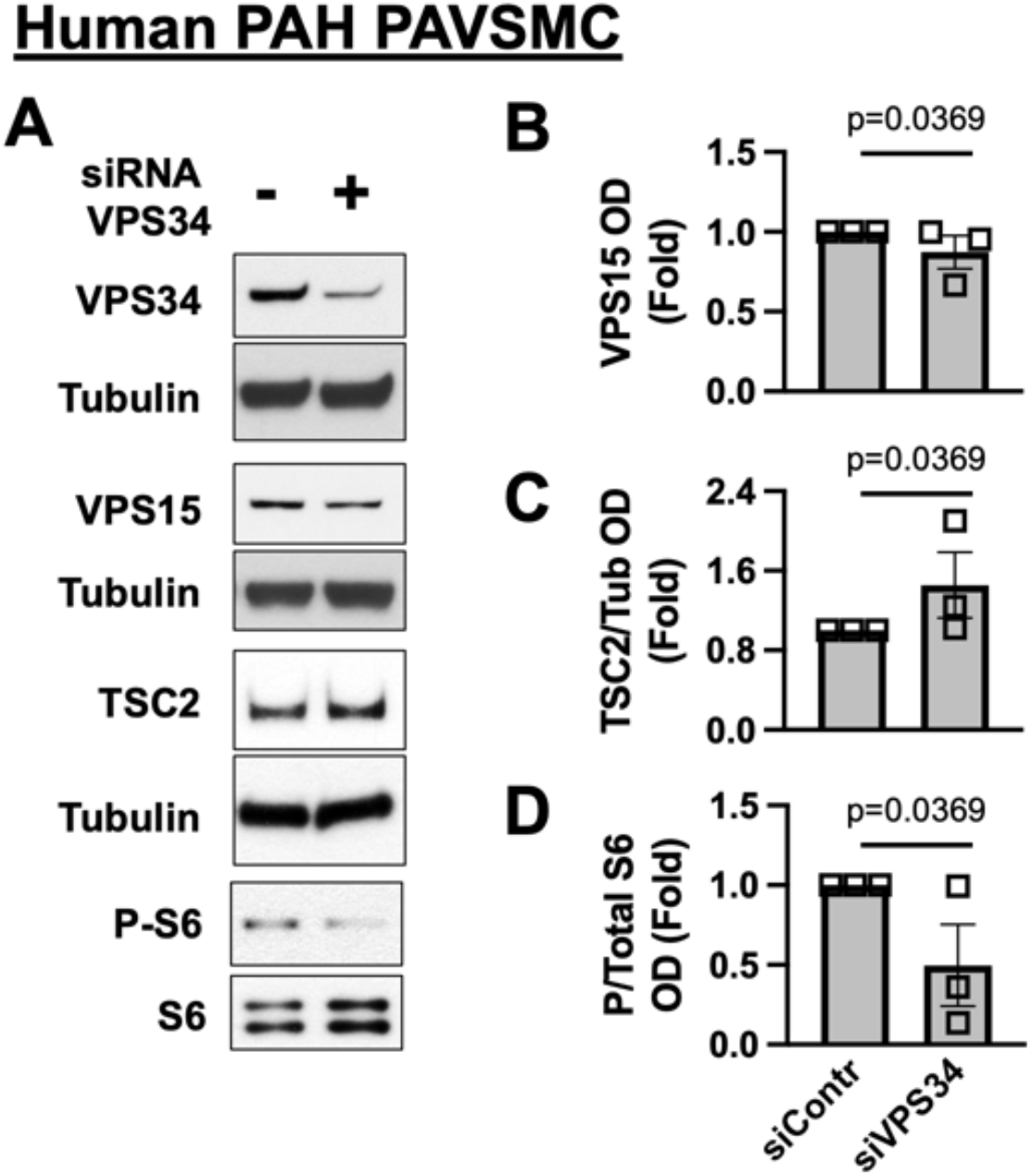
VPS34 regulates VPS15 and TSC2 in human PAH PAVSMCs. **A-D.**Human PAH PAVSMCs were transfected with siRNA VPS34 (+) or control siRNA (-); 72 hours later, immunoblot analysis was performed. Data are means±SE, fold change to control siRNA from n=3 subjects/group, by Mann-Whitney U test.

## Discussion

This study demonstrates that VPS34 activation, driven by the loss of inhibitory Ser164 phosphorylation (P-S164-VPS34), plays a pivotal role in sustaining the hyperproliferative phenotype of PAVSMCs and driving pulmonary vascular remodeling in PAH. Furthermore, we establish a novel role for activated VPS34 in mediating PAVSMC-derived FN production and secretion, linking cell proliferation to ECM remodeling. Mechanistically, our data suggest an upstream regulatory axis wherein PPPs, such as PPP1CA, coordinate with Akt to mediate the dephosphorylation and subsequent activation of VPS34. Downstream, active VPS34 suppresses the mTOR negative regulator TSC2, fueling S6 phosphorylation and cell proliferation. Finally, pharmacological inhibition of VPS34 via SAR405 successfully blunted PAVSMC proliferation *in vitro* and attenuated experimental pulmonary vascular remodeling *in vivo*, identifying the VSM VPS34 axis as a promising therapeutic target in PAH (**Figure 6**).

**Figure 6.**
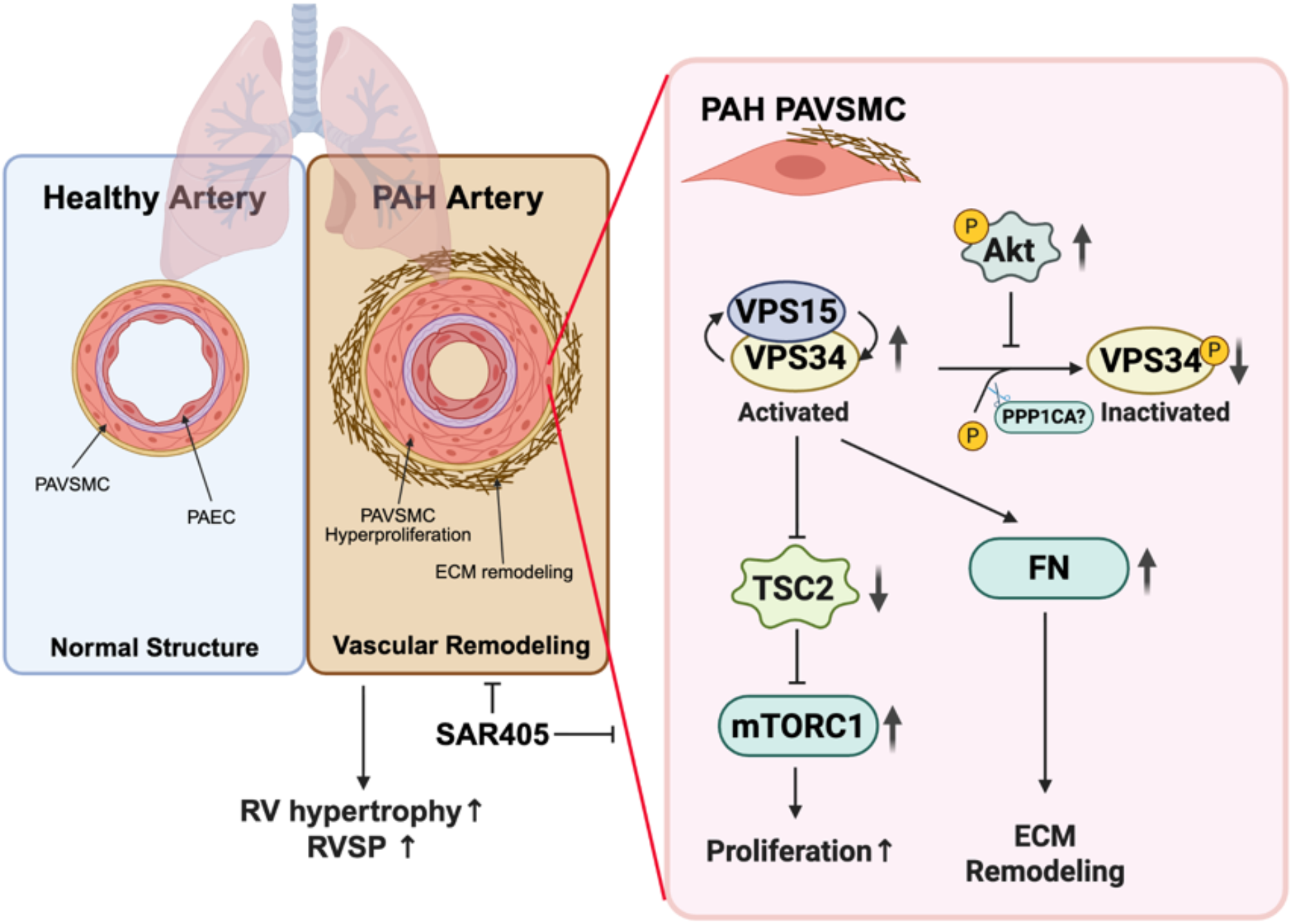
Schematic representation of the mechanism by which increased activation of VPS34 in PAVSMCs promotes pulmonary vascular remodeling and PH. In PAH PAVSMCs, increased Akt activation drives VPS34 activation via loss of inhibitory S164 phosphorylation. This is possibly mediated through PPPs, such as PPP1CA. Active VPS34 downregulates TSC2, activating mTORC1, to promote PAVSMC hyperproliferation, together with an increase in FN production. The selective VPS34 inhibitor SAR405 attenuates the pathogenic changes in PAH PAVSMC and male mice exposed to SuHx, highlighting the VPS34 axis as a therapeutic target in PAH. Figure generated using BioRender. Abbreviations: PAVSMC – pulmonary arterial vascular smooth muscle cell; PAEC – pulmonary arterial endothelial cell; ECM – extracellular matrix; RV -right ventricle; RVSP – right ventricular systolic pressure; VPS – vacuolar protein sorting; PPP -protein phosphatase; TSC2 – tuberous sclerosis complex 2; mTORC1 – mammalian target of rapamycin complex 1; FN – fibronectin.

Our findings establish that the loss of the P-S164-VPS34 inhibitory brake drives PAVSMC hyperproliferation and pulmonary vascular remodeling. Both siRNA-mediated knockdown of VPS34 (**Figure 1 E,F**) and direct enzymatic inhibition using the selective small-molecule inhibitor SAR405 (**Figure 1 G,H**) significantly reduced the proliferation marker Ki67 in human PAH PAVSMCs. This pathobiological relevance was further validated *in vivo*, where a pronounced decrease in P-S164-VPS34 closely correlated with the severity of pulmonary vascular remodeling in SuHx mice (**Figure 2 E-G**).

However, the precise upstream mechanisms dictating VPS34 dephosphorylation in the pulmonary vasculature warrant deeper exploration. We show for the first time that Akt is intrinsically involved in regulating this specific VPS34 PTM in human PAVSMCs; suppressing Akt activity restored the inhibitory P-S164-VPS34 mark (**Figure 3 A,B**). Because Akt operates primarily as a kinase, its capacity to promote the dephosphorylated, activated state of VPS34 likely relies on intermediate phosphatases. Our *in silico* prioritization identified PPP1CA as a top candidate, ranking third among 13,045 nominated proteins (**Figure 3C-E**). While STRING-db aggregates and predicts PPIs largely based on documented experimental evidence and literature, it remains limited in predicting novel interactions that lack prior experimental support. Conversely, SPP nominates candidates from primary sequence and network/functional proximity rather than curated interaction evidence, though downstream bench-based validation remains essential. For Akt-PPP pairs, where most interactions are already documented, STRING-db and SPP produced concordant rankings (**Figure 3D**); because these pairs are well-characterized, this agreement is best read as a sanity check on the pipeline’s behavior on known biology rather than external validation of its accuracy on novel pairs. For VPS34-PPP pairs, where STRING-db returns little signal due to sparse prior literature, we used the SPP ranking to prioritize PPP1CA for experimental follow-up. Given prior evidence that PPP1CA contributes to uncontrolled cellular proliferation^25,26^ and physically interacts with Akt^27,28^, we hypothesize that Akt activation in PAH may scaffold or activate a PPP1CA-containing phosphatase complex to selectively strip the inhibitory phosphate from VPS34. We understand that VPS34 is expressed not only in PAVSMCs. Interestingly, our preliminary findings also reveal modest VPS34 activation within the endothelial layer of small pulmonary arteries (**Supplemental Figure 1**). Whether this endothelial VPS34 activation coordinates with the Akt axis to promote the characteristic apoptosis-resistant phenotype of PAH pulmonary arterial endothelial cells remains an intriguing avenue for future study.

Vascular remodeling in PAH is fundamentally coupled with extensive ECM deposition, and strategies aimed at arresting matrix deposition have shown profound efficacy in preclinical models. To our knowledge, this study is the first to demonstrate that VPS34 directly regulates FN production within PAVSMCs (**Figure 4**). This aligns with emerging paradigms in oncology and fibrosis, where VPS34 governs matrix dynamics; for instance, LLGL2-dependent VPS34 regulation modulates FN expression in prostate cancer cells^29^, and its inhibition via PIK-III diminishes FN deposition in activated fibroblasts^30^. Our data expand upon this biology, confirming that targeting VPS34 with SAR405 downregulates PAVSMC-derived FN both *in vitro* and within the remodeled medial layer *in vivo*.

Beyond matrix production, active VPS34 extensively rewires intracellular growth signaling. Class III PI3Ks typically assemble into distinct functional complexes structured around the stabilizing protein VPS15^6,31^. Our observation that VPS34 silencing concomitantly reduces VPS15 levels suggests a mechanism of mutual protein stabilization within PAVSMCs (**Figure 5B**). Concurrently, we found that VPS34 activation suppresses TSC2 (**Figure 5C**), a critical negative regulator of mTORC1 known to be downregulated in PAH^4^. In fibroblast models, VPS34 can interact with TSC1, disrupting the TSC1/2 heterodimer and targeting TSC2 for degradation^11^. Our data suggest a similar disruptive mechanism may operate in hyperproliferative PAVSMCs. By downregulating TSC2, activated VPS34 increases downstream S6 phosphorylation (**Figure 5D**), effectively coupling endosomal trafficking or autophagic signaling directly to the metabolic engine of the mTOR pathway to drive vascular remodeling^6^.

To verify the consistency of our experimental conditions with published data^4^, we analyzed newly received human PAVSMCs from PHBI. The IB analysis showed a trend of an increase in FN and a decrease in TSC2 (**Supplemental Figure 2**). However, we acknowledge several limitations in the current study. First, while P-S164-VPS34 serves as a robust surrogate for kinase suppression, direct *in vitro* VPS34 lipid kinase activity assays will be valuable to fully define its enzymatic kinetics in diseased tissue. Second, our *in vivo* therapeutic evaluation was restricted to male mice; given the established sex paradox in PAH^32,33^, validation in female mice and alternative preclinical models (such as monocrotaline- or SuHx-treated rats) is essential. Third, while SAR405 demonstrated excellent efficacy against SuHx-induced pulmonary vascular remodeling, small-molecule inhibitors inherently carry risks of off-target effects, and SAR405 itself is not currently clinically approved. Nonetheless, our proof-of-concept genetic and pharmacological data strongly justify the future development of highly selective, bioavailable VPS34 inhibitors tailored for pulmonary vascular diseases.

For decades, the clinical management of PAH has relied predominantly on vasodilators targeting the endothelin, nitric oxide, and prostacyclin pathways^34^. While these agents improve hemodynamics and quality of life, and have some antiproliferative effects, they fail to arrest or reverse the underlying occlusive vascular remodeling. The recent clinical approval of the activin signaling inhibitor, sotatercept, represents a paradigm shift, establishing that therapies directly targeting vascular remodeling can yield profound clinical benefits in PAH, with expanding potential in other WHO PH Groups. By demonstrating that VPS34 inhibition simultaneously blunts PAVSMC hyperproliferation and halts pathogenic fibronectin deposition, our study positions the VPS34 axis as a compelling, multi-pronged therapeutic target capable of addressing the root proliferative and fibrotic drivers of pulmonary hypertension.

## Abbreviations

PAH: Pulmonary arterial hypertension
PAVSMC: Pulmonary arterial vascular smooth muscle cell
VSM: Vascular smooth muscle
PI3K: Phosphatidylinositol-3 kinase
SUMO: Small ubiquitin-like modifier
PTM: Posttranslational modification
TSC2: Tuberous sclerosis complex 2
VPS: Vacuolar protein sorting
mTOR: Mammalian target of rapamycin
PtdIns3P: Phosphatidylinositol 3-phosphate
RVSP: Right ventricular systolic pressure
PA: MTPulmonary arterial medial thickness
PPP: Protein phosphatase

## Acknowledgments

None.

## Figure legends

**Supplemental Figure 1.**

**A,B:** IHC analysis of human lung tissues to detect P-S164-VPS34 (red), CD31/VWF (green), and DAPI (blue). Bar=30 μm. Representative images (**A**) and optical density (OD) measurements (**B**) of P-S164-VPS34 in CD31/vWF-positive regions of human small pulmonary arteries. Data are means±SE from n=4 subjects/group, analyzed by Mann-Whitney U test.

**Supplemental Figure 2.**

Immunoblot analysis confirming increased FN and decreased TSC2 in human PAH PAVSMC.

**Supplemental Table 1.**

**Information about the de-identified cell donors.**

## References

1 Mocumbi A, Humbert M, Saxena A, Jing Z-C, Sliwa K, Thienemann F, et al. Pulmonary hypertension. Nat Rev Dis Primers 2024;10:1–20. 10.1038/s41572-023-00486-7.

2 Goncharova EA, Ammit AJ, Irani C, Carroll RG, Eszterhas AJ, Panettieri RA, et al. PI3K is required for proliferation and migration of human pulmonary vascular smooth muscle cells. American Journal of Physiology-Lung Cellular and Molecular Physiology 2002;283:L354–63.

3 Berghausen EM, Janssen W, Vantler M, Gnatzy-Feik LL, Krause M, Behringer A, et al. Disrupted PI3K subunit p110α signaling protects against pulmonary hypertension and reverses established disease in rodents. The Journal of Clinical Investigation 2021;131:. 10.1172/jci136939.

4 Shen Y, Goncharov DA, Pena A, Baust J, Chavez Barragan A, Ray A, et al. Cross-talk between TSC2 and the Extracellular Matrix Controls Pulmonary Vascular Proliferation and Pulmonary Hypertension. Sci Signal 2022;15:eabn2743. 10.1126/scisignal.abn2743.

5 Bilanges B, Posor Y, Vanhaesebroeck B. PI3K isoforms in cell signalling and vesicle trafficking. Nat Rev Mol Cell Biol 2019;20:515–34. 10.1038/s41580-019-0129-z.

6 Shen Y, Gleghorn JP. Class III Phosphatidylinositol-3 Kinase/Vacuolar Protein Sorting 34 in Cardiovascular Health and Disease. J Cardiovasc Transl Res 2025. 10.1007/s12265-024-10581-z.

7 Stjepanovic G, Baskaran S, Lin MG, Hurley JH. Vps34 Kinase Domain Dynamics Regulate the Autophagic PI 3-Kinase Complex. Mol Cell 2017;67:528–534.e3. 10.1016/j.molcel.2017.07.003.

8 Gstrein T, Edwards A, Přistoupilová A, Leca I, Breuss M, Pilat-Carotta S, et al. Mutations in Vps15 perturb neuronal migration in mice and are associated with neurodevelopmental disease in humans. Nat Neurosci 2018;21:207–17. 10.1038/s41593-017-0053-5.

9 Su H, Liu W. PIK3C3/VPS34 control by acetylation. Autophagy 2018;14:1086–7. 10.1080/15548627.2017.1385676.

10 Yoon MS. Vps34 and PLD1 take center stage in nutrient signaling: their dual roles in regulating autophagy. Cell Commun Signal 2015;13:44. 10.1186/s12964-015-0122-x.

11 Mohan N, Shen Y, Dokmanovic M, Endo Y, Hirsch DS, Wu WJ. VPS34 regulates TSC1/TSC2 heterodimer to mediate RheB and mTORC1/S6K1 activation and cellular transformation. Oncotarget 2016;7:52239–54. 10.18632/oncotarget.10469.

12 Wang W, Li J, Tan J, Wang M, Yang J, Zhang ZM, et al. Endonuclease G promotes autophagy by suppressing mTOR signaling and activating the DNA damage response. Nat Commun 2021;12:476. 10.1038/s41467-020-20780-2.

13 Hong-Brown LQ, Brown CR, Navaratnarajah M, Lang CH. FoxO1-AMPK-ULK1 Regulates Ethanol-Induced Autophagy in Muscle by Enhanced ATG14 Association with the BECN1-PIK3C3 Complex. Alcohol Clin Exp Res 2017;41:895–910. 10.1111/acer.13377.

14 Furuya T, Kim M, Lipinski M, Li J, Kim D, Lu T, et al. Negative regulation of Vps34 by Cdk mediated phosphorylation. Mol Cell 2010;38:500–11. 10.1016/j.molcel.2010.05.009.

15 Kumar B, Ahmad R, Sharma S, Gowrikumar S, Primeaux M, Rana S, et al. PIK3C3 Inhibition Promotes Sensitivity to Colon Cancer Therapy by Inhibiting Cancer Stem Cells. Cancers (Basel) 2021;13:. 10.3390/cancers13092168.

16 Jiang X, Bao Y, Liu H, Kou X, Zhang Z, Sun F, et al. VPS34 stimulation of p62 phosphorylation for cancer progression. Oncogene 2017;36:6850–62. 10.1038/onc.2017.295.

17 Yao Y, Li H, Da X, He Z, Tang B, Li Y, et al. SUMOylation of Vps34 by SUMO1 promotes phenotypic switching of vascular smooth muscle cells by activating autophagy in pulmonary arterial hypertension. Pulm Pharmacol Ther 2019;55:38–49. 10.1016/j.pupt.2019.01.007.

18 Kudryashova TV, Goncharov DA, Pena A, Kelly N, Vanderpool R, Baust J, et al. HIPPO– Integrin-linked Kinase Cross-Talk Controls Self-Sustaining Proliferation and Survival in Pulmonary Hypertension. Am J Respir Crit Care Med 2016;194:866–77. 10.1164/rccm.201510-2003OC.

19 Kim J, Kim YC, Fang C, Russell RC, Kim JH, Fan W, et al. Differential regulation of distinct Vps34 complexes by AMPK in nutrient stress and autophagy. Cell 2013;152:290–303. 10.1016/j.cell.2012.12.016.

20 Su H, Yang F, Wang Q, Shen Q, Huang J, Peng C, et al. VPS34 Acetylation Controls Its Lipid Kinase Activity and the Initiation of Canonical and Non-canonical Autophagy. Mol Cell 2017;67:907–921.e7. 10.1016/j.molcel.2017.07.024.

21 Xie W, Jin S, Wu Y, Xian H, Tian S, Liu DA, et al. Auto-ubiquitination of NEDD4-1 Recruits USP13 to Facilitate Autophagy through Deubiquitinating VPS34. Cell Reports 2020;30:2807–2819.e4. 10.1016/j.celrep.2020.01.088.

22 Jiang L, Goncharov DA, Shen Y, Lin D, Chang B, Pena A, et al. Akt-Dependent Glycolysis-Driven Lipogenesis Supports Proliferation and Survival of Human Pulmonary Arterial Smooth Muscle Cells in Pulmonary Hypertension. Front Med (Lausanne) 2022;9:886868. 10.3389/fmed.2022.886868.

23 Fresno Vara JA, Casado E, de Castro J, Cejas P, Belda-Iniesta C, González-Barón M. PI3K/Akt signalling pathway and cancer. Cancer Treat Rev 2004;30:193–204. 10.1016/j.ctrv.2003.07.007.

24 Zuo W, Liu N, Zeng Y, Xiao Z, Wu K, Yang F, et al. Luteolin Ameliorates Experimental Pulmonary Arterial Hypertension via Suppressing Hippo-YAP/PI3K/AKT Signaling Pathway. Front Pharmacol 2021;12:663551. 10.3389/fphar.2021.663551.

25 Chen Z, Li Y, He K, Yang J, Deng Q, Chen Y, et al. CircGPRC5A enhances colorectal cancer progress by stabilizing PPP1CA and inducing YAP dephosphorylation. J Exp Clin Cancer Res 2023;42:334. 10.1186/s13046-023-02915-7.

26 Liu X, Xu G, Luo W, Wang K, Wang F. PPP1CA promotes hepatocellular carcinoma progression in a YAP1-dependent way. Cell Signal 2025;134:111938. 10.1016/j.cellsig.2025.111938.

27 Chen M, Wan L, Zhang J, Zhang J, Mendez L, Clohessy JG, et al. Deregulated PP1α phosphatase activity towards MAPK activation is antagonized by a tumor suppressive failsafe mechanism. Nat Commun 2018;9:159. 10.1038/s41467-017-02272-y.

28 Xiao L, Gong L-L, Yuan D, Deng M, Zeng X-M, Chen L-L, et al. Protein phosphatase-1 regulates Akt1 signal transduction pathway to control gene expression, cell survival and differentiation. Cell Death Differ 2010;17:1448–62. 10.1038/cdd.2010.16.

29 Hong G-L, Kim K-H, Kim Y-J, Lee H-J, Cho S-P, Han S-Y, et al. Novel role of LLGL2 silencing in autophagy: reversing epithelial-mesenchymal transition in prostate cancer. Biol Res 2024;57:25. 10.1186/s40659-024-00499-w.

30 Sanchez S, McDowell-Sanchez AK, Al-Meerani SB, Cala-Garcia JD, Waich Cohen AR, Ochsner SA, et al. PIK-III exerts anti-fibrotic effects in activated fibroblasts by regulating p38 activation. PLoS One 2024;19:e0306624. 10.1371/journal.pone.0306624.

31 Yan Y, Flinn RJ, Wu H, Schnur RS, Backer JM. hVps15, but not Ca2+/CaM, is required for the activity and regulation of hVps34 in mammalian cells. Biochem J 2009;417:747–55. 10.1042/bj20081865.

32 Bousseau S, Gu S, Lahm T. Biological and Clinical Implications of Sex Differences in Right Heart Failure Associated With Pulmonary Arterial Hypertension. Arterioscler Thromb Vasc Biol 2026;46:e322096. 10.1161/ATVBAHA.126.322096.

33 Shen Y, Goncharov DA, Avolio T, Ray A, Okorie E, DeLisser H, et al. Differential effects of integrin-linked kinase inhibitor Cpd22 on severe pulmonary hypertension in male and female rats. Pulm Circ 2020;10:2045894019898593. 10.1177/2045894019898593.

34 Humbert M, Kovacs G, Hoeper MM, Badagliacca R, Berger RMF, Brida M, et al. 2022 ESC/ERS Guidelines for the diagnosis and treatment of pulmonary hypertension. Eur Heart J 2022;43:3618–731. 10.1093/eurheartj/ehac237.

35 Gonyea CR, Shen Y, Nelson KM, Bird RN, Gilbert RM, Olutoye OO, et al. The nitrofen/bisdiamine murine model of congenital diaphragmatic hernia has a pulmonary hypertension vascular phenotype consistent with human CDH. Am J Physiol Lung Cell Mol Physiol 2025;329:L48–60. 10.1152/ajplung.00233.2024.

36 Kelly NJ, Dandachi N, Goncharov DA, Pena AZ, Radder JE, Gregory AD, et al. Automated Measurement of Blood Vessels in Tissues from Microscopy Images. Curr Protoc Cytom 2016;78:12.44.1-12.44.13. 10.1002/cpcy.10.

