## Supplemental materials for "VPS34 regulation in pulmonary arterial vascular smooth muscle cells in pulmonary hypertension"

**Methods**

**Immunohistochemical, Immunocytochemical, and Immunoblot Analyses**

These analyses were performed as previously described[4,18,35]. Immunohistochemical (IHC) and immunocytochemical (ICC) images were captured with the All-in-One Fluorescence Microscope BA-X810 (Keyence), Leica TCS SP8 Confocal Microscope (Leica), or Olympus BX61VS Slide Scanner (Olympus). Immunoblot (IB) signals were captured with HyBlot CL Autoradiography Films (Thomas Scientific) and fixed and developed by the Medical Film Processor (Konica Minolta), or imaged by Azure Biosystems 400 (Azure). Optical density measurements for IHC, ICC, and IB were done with ImageJ (NIH). Antibodies for Tubulin (#2148), VPS34 (PIK3C3, #4263), Akt (#9272), P-S473-Akt (#4060), TSC2 (#4308), VPS15 (#14580), Ki-67(#9129), S6 (#2217), P-Ser235/236-S6 (#4856), and HRP-linked anti-rabbit IgG (#7074), were purchased from Cell Signaling (Boston, MA). Antibodies for P-Ser164-VPS34 (SAB1305578) and α-Smooth Muscle Actin (SMA)-FITC (F3777) were purchased from Sigma-Aldrich (Burlington, MA). Antibodies for fibronectin (Ab2413) and Von Willebrand Factor (Ab11713) were purchased from AbCam (Waltham, MA). Anti-PPP1CA (A12468) was purchased from ABclonal (Woburn, MA). Anti-CD31 (AF3628) was purchased from R&D Systems BioTechne (Devens, MA). Donkey anti-Goat IgG (H+L) Highly Cross-Adsorbed Secondary Antibody, Alexa Fluor™ Plus 488 (A32814TR), and goat anti-rabbit IgG (H+L) Cross-Adsorbed Secondary Antibody, Alexa Fluor™ 594 (A11012) were purchased from Invitrogen Thermo Fisher Scientific (Waltham, MA). Anti-IgG Donkey Antibody, Alexa Fluor® 488 (VWR Catalog #102649-960) was purchased from Jackson ImmunoResearch Labs (West Grove, PA). The dilutions of the antibodies for IHC, ICC, and IB analyses were prepared according to the manufacturer's instructions.

**Animals**

Six- to eight-week-old C57BL/6J mice were randomly assigned to experimental groups and exposed to hypoxia (10% $O_2$) for up to 28 days. Subcutaneous injections of SU5416 (20mg/kg) (BioTechne) were performed on days 0, 7, and 14 of the experiment. On Day 15, mice exposed to SuHx, while still maintained under hypoxic conditions, were randomly assigned to two groups. SAR405 (MedChemExpress), suspended in warm corn oil (ACROS Organics) (10 mg/kg/day, oral gavage, 200 μL/mouse), or warm corn oil alone (Veh, oral gavage, 200 μL/mouse) was administered to the randomly assigned groups of mice daily for 14 days. Controls were mice maintained under normoxia.

Blinded hemodynamic analysis was performed as previously described[4,18,33]. Briefly, animals were anesthetized with isoflurane: 5% for induction, 2% during surgery, and 1% while performing Pressure-Volume (PV) loop measurements (Kent Scientific). PV loop measurements were performed using PV catheters (Sciensence). The catheter attached to the data acquisition system (AdInstruments) was inserted into the RV. Data were acquired by the MPVS ULTRA (Millar, Pearland, TX) and PowerLab 8/35 (AdInstruments, Colorado Springs, CO), and were analyzed by LabChart8 (AdInstruments)

Blinded morphological and immunohistological analyses were performed as described previously[4,18,33]. Briefly, after being perfused with ice-cold PBS through the RV, the lungs were fixed in 4% paraformaldehyde in PBS and embedded in paraffin. After sectioning, lung tissue slides (5 μm thickness) were stained with H&E or immunostained to detect P-Ser164-VPS34, fibronectin, SMA, vWF, CD31, and nuclei (DAPI) as previously described[4,35]. Images for PA medial thickness (PA MT) calculations were captured from blindly selected small PAs (around 50 μm outer diameter from a minimum of six animals/groups and a minimum of 12 PAs/animal) of H&E-stained lung tissues, and calculated using the VMI calculator[4,18,36].

**Bioinformatics**

Disease context was supplied as the twelve PAH genes with definitive evidence in ClinGen: BMPR2, ACVRL1, ATP13A3, CAV1, EIF2AK4, ENG, GDF2, KCNK3, KDR, SMAD9, SOX17, and TBX4. We included them because we assumed the intermediate protein of interest was not just any phosphatase but one plausibly involved in the disease, so both their annotations and their interaction profiles were scored. This distinguished a candidate sitting close to the disease proteins from one connected only to the anchors.

Interactions were scored as a complete square over the anchors, the disease proteins, and the whole proteome rather than an anchor-against-proteome strip, because a promiscuous protein scores highly against everything and only a proteome-wide background separates a specific partner from a hub. Each pair was therefore judged on how far it exceeded what both partners' promiscuity alone predicted, correcting for the anchor's promiscuity as well as the candidate's. Two-hop candidates were admitted at a discount, carrying the weaker edge of their best path and the intermediate that found them.

Candidates were required to share a subcellular compartment with the anchors, which resolved to cytoplasm, membrane, mitochondria, nucleus, and peroxisome. The phosphatase requirement was pinned to four identifiers GO:0004721, GO:0004722, GO:0006470, and EC 3.1.3.16. The disease requirement combined ten specified terms covering BMP signalling, angiogenesis, hypoxia response, endothelial proliferation, and potassium transport with 128 terms shared by at least two of the twelve disease genes. Terms were down-weighted by how common they were across the proteome, so generic annotation did not carry the ranking.

In total, all 20,645 proteins were scored against the anchors, and 13,045 proteins matching at least one mechanism term were nominated and ranked. Candidate scores nominally combined standardized interaction and mechanism axes equally; the mechanism axis combined functional annotation with disease annotation and disease proximity. The interaction axis corrected predicted PPI probabilities for both partners' proteome-wide baseline interaction rates, and co-localization was used as a gating criterion.

The output was used to prioritize candidates for experimental testing. We then used the PPI predictions alongside STRING-db to further explore predicted interactions among protein phosphatases, VPS34, and Akt.
