## Supplemental figures for "VPS34 regulation in pulmonary arterial vascular smooth muscle cells in pulmonary hypertension"

**Supplemental Table 1**

| Biobank # | Age | Sex | Donor Condition |
|---|---|---|---|
| 5414 | 40 | Male | Healthy |
| 4914 | 33 | Female | Healthy |
| 1515 | 19 | Male | Healthy |
| 109 | 50 | Female | Healthy |
| 125 | 37 | Female | Healthy |
| L116 | 57 | Female | Healthy |
| L136 | 47 | Female | Healthy |
| L104 | 25 | Male | Healthy |
| ki | 40 | Female | IPAH |
| 7 | 62 | Female | IPAH |
| 25 | 39 | Female | IPAH |
| 21414 | 31 | Male | IPAH |
| 11 | 58 | Female | IPAH |
| 157 | 22 | Female | IPAH |
| 10816 | 53 | Female | PAH Scleroderma |
| L85 | 56 | Female | IPAH |
| L95 | 40 | Male | IPAH |
| L10 | 47 | Female | IPAH |
| L13 | 15 | Female | IPAH |

**Supplemental Figure 1**

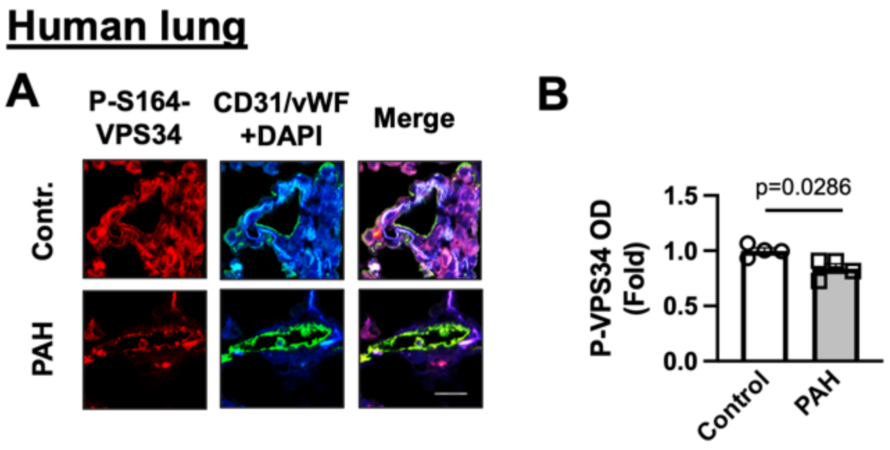

**Supplemental Figure 2**

### Human PAVSMC

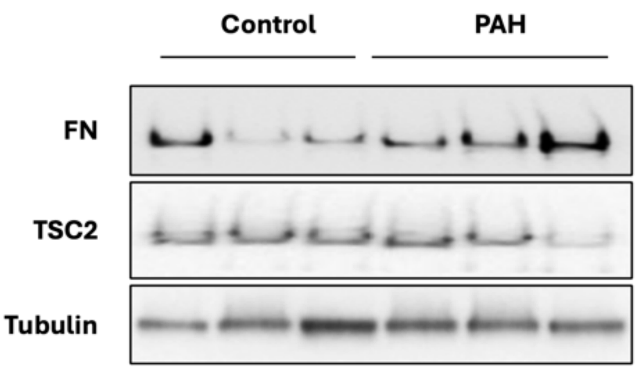
